# Multiple roles of PIWIL1 in mouse neocorticogenesis

**DOI:** 10.1101/106070

**Authors:** Barbara Viljetic, Liyang Diao, Jixia Liu, Zeljka Krsnik, Sagara H.R. Wijeratne, Ryan Kristopovich, Marina Dutre-Clarke, Matthew L. Kraushar, Jimin Song, Jinchuan Xing, Kevin C. Chen, Mladen-Roko Rasin

## Abstract

PIWI-interacting RNAs (piRNAs) and their associated PIWI proteins play an important role in repressing transposable elements in animal germlines. However, little is known about the function of PIWI proteins and piRNAs in the developing brain. Here, we investigated the role of an important PIWI family member, Piwi-like protein 1 (Piwil1; also known as Miwi in mouse) in the developing mouse neocortex. Using a *Piwil1* knock-out (*Piwil1* KO) mouse strain, we found that Piwil1 is essential for several steps of neocorticogenesis, including neocortical cell cycle, neuron migration and dendritogenesis. Piwil1 deletion resulted in increased cell cycle re-entry at embryonic day 17 (E17) when predominantly intracortically projecting neurons are being produced. Prenatal Piwil1 deletion increased the number of Pax6+ radial glia at postnatal day 0 (P0). Furthermore, Piwil1 deletion disrupted migration of Satb2+ neurons within deep layers at E17, P0 and P7. Satb2+ neurons showed increased co-localization with Bcl11b (also known as Ctip2), marker of subcortically projecting neurons. Piwil1 knockouts had disrupted neocortical circuitry represented by thinning of the corpus callosum and altered dendritogenesis. We further investigated if Piwil1 deletion disrupted expression levels of neocortical piRNAs by small RNA-sequencing in neocortex. We did not find differential expression of piRNAs in the neocortices of *Piwil1* KO, while differences were observed in other *Piwil1* KO tissues. This result suggests that Piwil1 may act independently of piRNAs and have novel roles in higher cognitive centers, such as neocortex. In addition, we report a screen of piRNAs derived from tRNA fragments in developing neocortices. Our result is the first report of selective subsets of piRNAs and tRNA fragments in developing prenatal neocortices and helps clarify some outstanding questions about the role of the piRNA pathway in the brain.

## Introduction

The neocortex is vital for major high-level functions, including voluntary motor commands and consciousness. The mature six-layered neocortex develops in a precisely timed and highly regulated manner (1–5). All projection neurons of the neocortex arise from a pseudolayered ventricular zone containing specialized neural stem cells called radial glia (RG). RG undergo a cell cycle like other neural stem cells, but the dynamics of the cell cycle change as neocortical neurogenesis progresses (6,7). During neocortical neurogenesis, RG divide in the ventricular zone (VZ) to become either multipotent intermediate progenitor cells (IPC) or postmitotic neuroblasts. Neuroblasts then migrate out of VZ and populate the cortical plate that will ultimately consist of six functionally distinct layers (6–9). Here, deep layer neurons will predominantly express distinct transcription factor identity markers like B-Cell CLL/Lymphoma 11B (Bcl11b; also known as Ctip2) and Transducin-Like Enhancer of Split 4 (Tle4), and will predominantly project subcortically to innervate the thalamus, brain stem and spinal cord. The upper layer neurons will migrate out of VZ into Cortical plate (CP), surpass the deep layers to reside in superficial layers completing the "inside-out" fashion of neocortical formation by early postnatal days. The upper layer neurons will express distinct transcription factors like Cut-Like Homeobox 1 (*Cutl1, Cdp*) and Special AT-rich sequence-binding protein 2 (Satb2). However, Satb2 was recently found in subcortically projection neurons too (10,11). The later-born neurons will only project intracortically, with some forming the corpus callosum, the largest commisure connecting the hemispheres and whose formation is disrupted in numerous neurodevelopmental disorders (12,13). Small aberrations in neocortical development can have devastating life-long consequences and, thus, the underpinning molecular mechanisms are widely studied.

Recently, using RNAi knock-down experiments, Piwi-Like RNA-Mediated Gene Silencing 1 (Piwil1) was described to contribute to neocortical development in rodents (14). In addition, a subset of autism patients has *de novo* mutations in two PIWI genes (15). However, the role of Piwi-interacting RNAs (piRNAs) in these phenotypes and their expression in developing neocortices is still unclear. piRNAs are 26-30nt long non-coding RNAs that are related to but clearly distinct from microRNAs (16). They are predominantly found in animal germlines where their best understood function is to repress transposable elements by guiding PIWI proteins, members of the Argonaute protein family, to their targets (16). The piRNA pathway also has a role in memory formation in Aplysia neurons (17), while piRNAs were found to be expressed in the mouse brain (18). piRNA associated PIWI proteins, such as Piwil1, have also been found to regulate transposon expression in the Drosophila brain (19). Since *Piwil1* mRNA is expressed in the developing neocortical wall and increases from early (E13) to late neurogenesis (E15 and E18), we hypothesized a role for Piwil1 in the later stages of neocortical neurogenesis and in neocortical circuit formation (5).

Here, we tested Piwil1 function in neocortical development by analyzing a *Piwil1* knock-out (KO) mouse at different developmental stages. We found that *Piwil1* is critical for normal cell cycle dynamics of RG, migration of later born neurons, and neocortical circuitry. To determine if piRNAs are involved in neocortical development, we investigated the expression of piRNAs in developing neocortices, spleen and testis of *Piwil1* KOs and littermate wild type (WT) mice using small RNA sequencing. Interestingly, in contrast to spleen and testis, neocortices depleted of *Piwil1* did not show significant differences in piRNA levels, indicating neocortex specific roles for Piwil1 that do not depend on piRNA levels. In addition, within the small RNA pool we identified a number of tRNA fragments that have piRNA characteristics. In recent years, there have been a number of reports of functional small RNAs in several species and cell-types derived from tRNA fragments, such as mammalian sperm. tRNA fragments were described to have miRNA like regulation of gene expression (20,43,44). In addition, these fragments were shown to regulate translation, likely targeting translational elongation (21). Our data suggest that tRNA fragments are present in the developing mouse neocortex and thus might play a role in neocortex development, perhaps through mRNA translation, a step in gene expression control that is becoming one of the key mechanisms in neocortical development (5,12,22,23).

## Material and methods

### Subjects

All mouse experiments were approved by and carried out in compliance with Rutgers University Robert Wood Johnson Medical School Institutional Animal Care and Use Committee protocols. *Piwil1* KO mice (purchased from JAX Mice, The Jackson laboratory) were bred as littermates from heterozygous parents and genotyped. For generation of embryonic *Piwil1* deletion mice, we performed timed pregnancies in which plugs found the next day were considered E0.5.

Adult mice were analyzed using a Golgi method for dendrites; while other analyses were performed at E13-E17, P0 or P7. All quantitative studies were run blind with respect to subject genotype.

### Immunohistochemistry

Mice embryonic brains were dissected and fixed by immersion in 4% paraformaldehyde (PFA) in phosphate-buffered saline (PBS), pH 7.4. Postnatal mouse brains were perfused with 4% PFA (pH 7.4), post-fixed overnight at 4°C and immersed in 30% sucrose in PBS. Brains were sectioned coronally using a vibratome (Leica) at 70 μm. Immunohistochemical experiments were performed as described previously (12,24). The following primary antibodies were used in dilution: 1:250 mouse anti-Sabt2 (catalog #ab51502, Abcam), 1:250 rat anti-Ctip2 (Bcl11b) (catalog #ab18465; Abcam), 1:250 rabbit anti-Cdp (M22 catalog #SC-13024; SCBT), 1:250 mouse anti-Tle4 (E10 catalog #SC-365406; SCBT), 1:250 rabbit anti-Pax6 (catalog #901301; BioLegend), 1:250 rabbit anti-Tbr2 (catalog # ab23345; Abcam), 1:500 rat anti-L1 (catalog # mab5272; Millipore), and incubated overnight at 4C. Appropriate secondary antibodies, raised in donkey, conjugated to cyanine 2 (Cy2), Cy3, Cy5 or Alexa were used at 1:250 dilution (Jackson ImmunoResearch Laboratories) for 2 hours at room temperature. Following DAPI staining, sections were mounted, coverslipped with Vectashield (Vector), and analyzed using an FV1000MPE microscope (Olympus) with 10 and 20x Olympus objectives.

### Cortical layer marker quantitative analysis

Confocal images of immunostained cerebral cortices of WT and Piwil1 KO (P0, P7) coronal sections were taken. The cortical image was then equally subdivided into columns of 10 virtual bins from cortical plate (bin 1) to ventricular zone (VZ) (bin 10). Analysis was performed by counting number of Cdp and Tle4-positive neurons or Bcl11b- and Satb2-positive neurons in each column of 10 bins (n=3 brains per age). Multivariate ANOVA (MANOVA) was conducted; if MANOVA was P < 0.05, then ANOVA was conducted. ANOVA was then conducted followed by post hoc analysis if ANOVA was P < 0.05. Post hoc analysis depended on the equality of variances. If Levene’s statistic proved that a given group had equal variances (P ≥ 0.05), then Tukey’s honest significant difference (HSD) was used for multiple comparison; for unequal variances (P < 0.05), Games–Howell was used. The standard error of the mean (SEM) is shown as the error bars in figures.

### Confocal microscopy and quantification

Images were taken with an Olympus BX61WI confocal microscope and processed using Fluoview FV-1000 software. Quantification of immunostaining and co-localization was done with Neurolucida and Image J software.

### Silencing of Piwil1 at E13 by in utero electroporation

Piwil1 expression was silenced in vivo by injecting *Piwil1* shRNA plasmid #4 along with Green Fluorescent Protein (GFP) at embryonic day 13 (E13) via *in utero* electroporation (IUE). Control shRNA with GFP was also injected. Embryos were harvested at E18 and coronal brain sections were analyzed using immunohistochemistry and confocal microscopy as described above, and quantification of distribution of GFP-positive neurons.

### Pulse-label thymidine analysis of in utero electroporation

To test cell cycle dynamics within the E17 neocortex, pulses of two thymidine analogues (CldU and IdU) were injected peritoneally at E16 (CldU, 24 hours prior to fixation) and E17 (IdU, 1 hour prior to fixation) as previously described (25). E17 embryo brains were fixed in 4% PFA overnight and sliced into 70 μm-thick sections. Processed tissue was immunostained for rat anti-BrdU (=anti-CldU, 1:100, Accurate), mouse anti-BrdU (=anti-IdU, 1:100, BD), rabbit anti-PH3 (phospho-histone H3, mitosis marker, 1:1000, Millipore) at E17 and analyzed using confocal microscopy. Double blind quantification was performed using automated software (MetaMorph).

### Golgi staining and analysis

Golgi staining in adult WT and *Piwil1* KO was performed on upper and lower layer neurons as per the manufacturer’s protocol (rapid Golgi kit; FD Neurotechnologies). Processed brains were sectioned in 240 micron slices and coverslipped as per the manufacturer’s protocol. Z-stack images (2um/step) were taken of at least 5 upper and 5 lower layer neocortical projection neurons from WT and KO (n=5 brains per age). Images were taken using a Leica DMRE bright-field microscope with Openlab software. Multi-tiff Z-stack images of neurons were reconstructed using off-site Neurolucida software and analyzed as described previously (26).

### PiRNA extraction and sequencing

Total RNA from seven neocortex samples (three WT and four KO), two testis samples (one WT and one KO) and one spleen (WT) sample were extracted using mirVana™ miRNA Isolation Kit (Ambion). Our previously published RNA size-selection and periodate oxidation and beta elimination protocol were used to specifically select for 2’ o-methylated small RNAs (27,28). We previously showed that this protocol produces very similar RNA profiles as Piwi protein immunoprecipitation protocols in both the human and Drosophila germline. However, we cannot exclude the possibility that there exist a novel species of small RNA in the mouse brain that is not a piRNA but is 2’ o-methylated. We also note that charged tRNAs are protected from degradation at the 3’ end, similar to 2’ o-methylation. The small RNA libraries were sequenced using Illumina HiSeq 2000 with 50 SE format at the Rutgers RUCDR Infinite Biologics Core Facility. The data has been deposited in SRA with accession number PRJNA309689.

### Bioinformatic analysis of the Illumina sequencing data

The Cutadapt software (version 1.4.1) was used to trim the sequencing reads and discarded any sequence reads of length <10nt. The read length distributions for all the brain samples (KO and WT) were very similar. The distribution for a representative sample (3WT) is presented in Supplementary figure 1. In all samples two major peaks exist, one around 17-20nts and one at 30nts.

**Figure 1.**
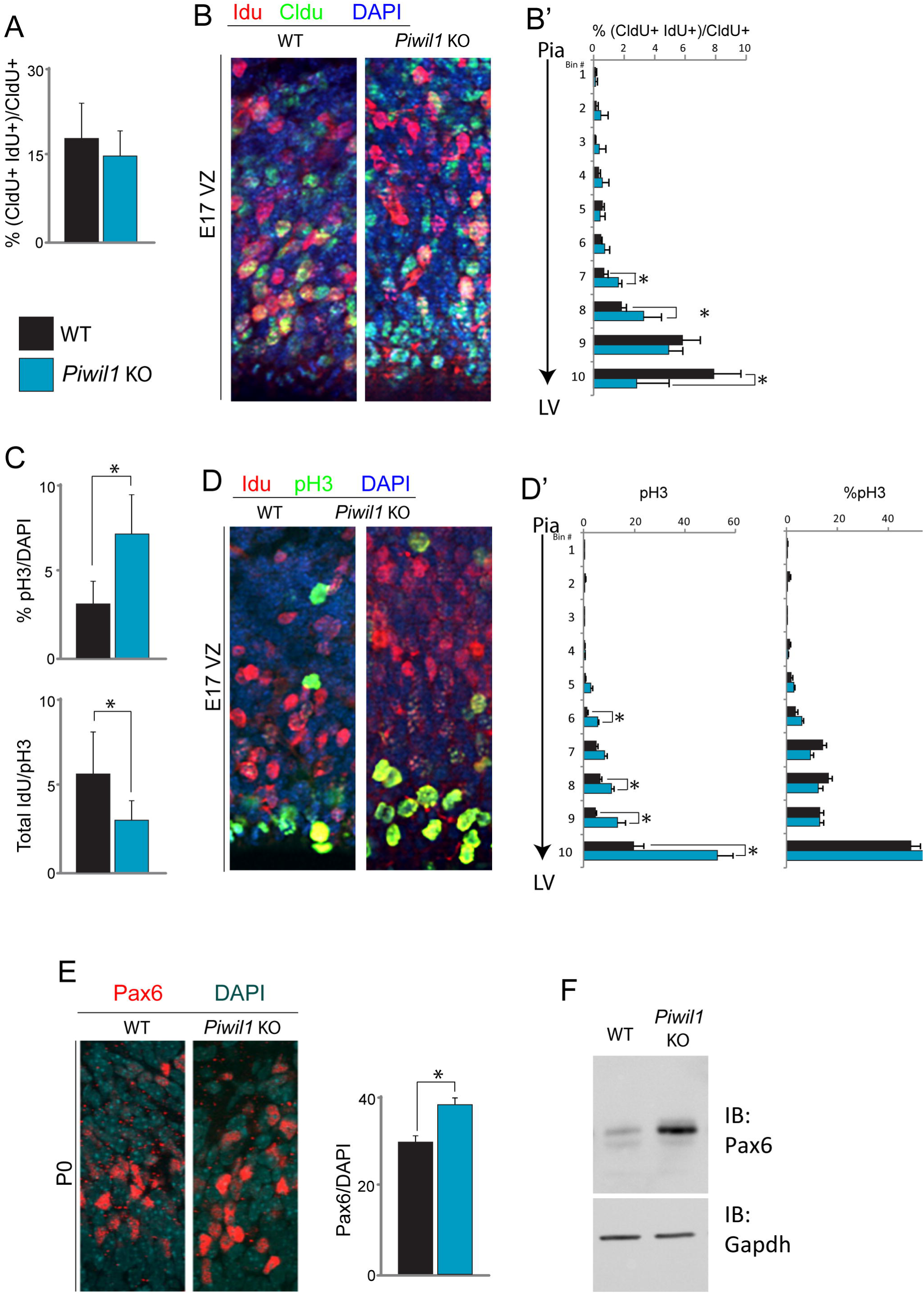
*Piwil1* knockout results in neocortical zone specific disruption of cell cycle. (A) Quantification of the proportion of CldU+ cells from total labeled cells (CldU and IdU) in developing neocortex of WT and *Piwil1* KO. (B) Representative confocal images of the developing VZ of WT and *Piwil1* KO at E17 labeled for tymidine analogues. Confocal images were divided into 10 equal bins for analysis from cortical plate (bin 1) to ventricular zone (bin 10). (B’) Quantification of the distribution of CldU+ (green) and IdU+ (red) cells in each bin is given as the percentage of the CldU+ cells from total labeled cells (CldU and IdU). DAPI is in blue. Mean bin proportion compared between WT and KO for each bin. (C) Quantification of the proportion of pH3+ cells from total DAPI labeled cells and proportion of IdU+ from pH3+ cells in neocortex of WT and *Piwil1* KO. (D) Confocal images of the WT and *Piwil1* KO VZ at E17 imunostained for IdU (red) and mytosis marker pH3 (green). (D’) Confocal images were divided into 10 equal bins as described above, and quantification of the distribution of pH3+ in each been is shown either as the percentage from total DAPI+ cells or as the comparison to IdU+. (E) Imunohistochemistry on P0 ventricular zone of WT and *Piwil1* KO for Pax6 radial glia progenitors (red). Quantification of the proportion of Pax6+ cells from total DAPI labeled cells showed increased number of Pax6+ in VZ of KO. (F) Western blot analysis of total neocortical lysate from WT and *Piwil1* KO at P0 for Pax6 (balanced to GAPDH).

All the reads were first aligned to a combined reference of non-coding RNAs (ncRNAs) annotated in Ensembl (GRCm38) and tRNAs annotated in the UCSC Genome Browser (mm10) using the Bowtie2 software (41). For this alignment step, both the ncRNA and tRNA reference sequences were buffered with Ns (i.e. characters that match any of the four nucleotides) on either end. This is because in many cases, we found that our reads aligned with the reference short RNA with an overhang of a few nucleotides (Supplementary figure 2), especially for reads aligning to tRNAs. Thus we used Bowtie 2 and allowed three ambiguous character matchings at each end of the sequence. Reads that aligned to any small RNAs in this step were removed before the remaining reads were aligned to the mouse genome (mm10) using Bowtie2 to identify piRNAs. All Bowtie2 alignments were performed using the following parameters unless otherwise noted. We allowed no mismatches and allowed 3 ambiguous character alignments (parameters: -p 8 -N 0 --n-ceil L,0,0.2).

**Figure 2.**
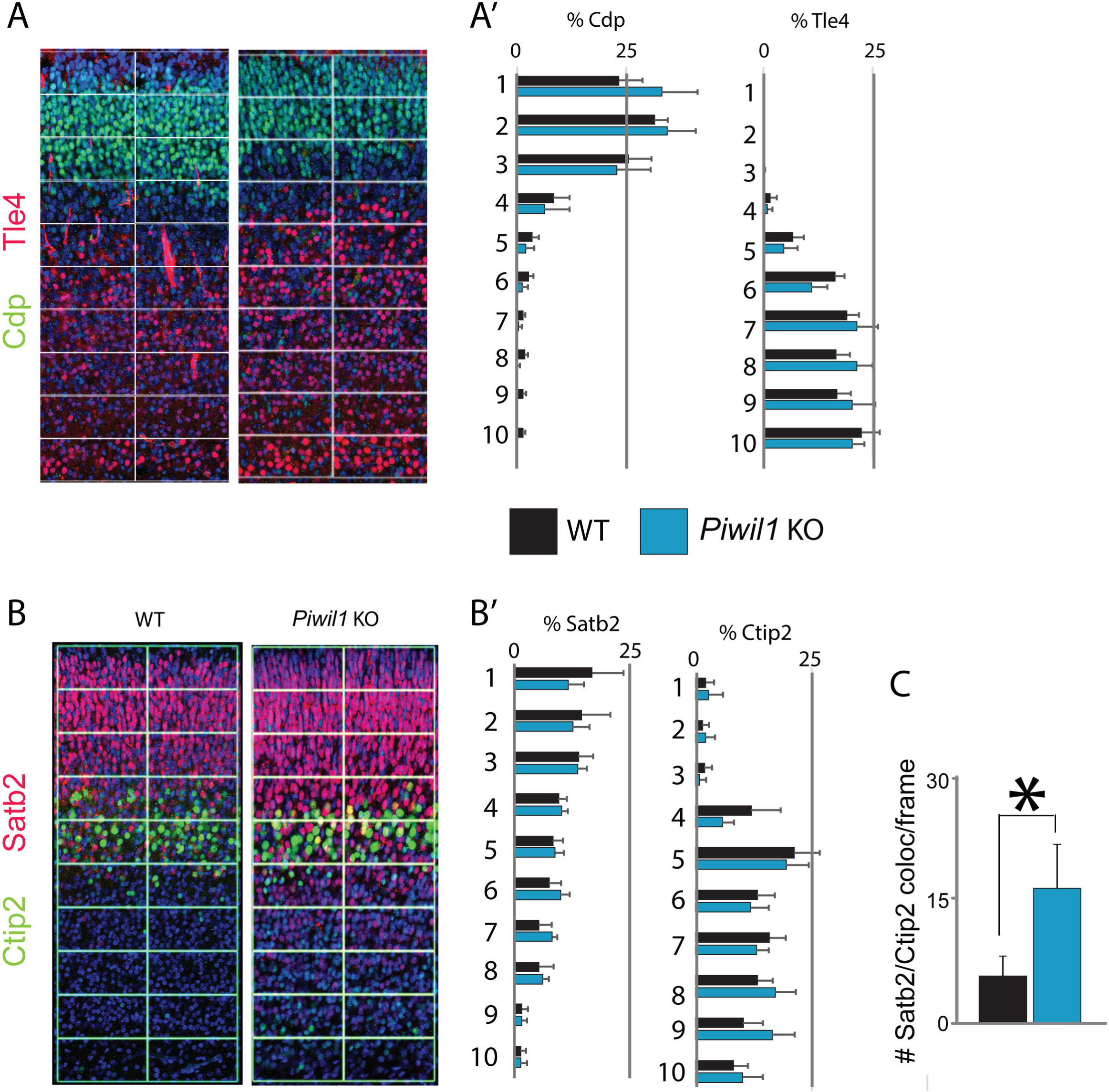
Distribution of subpopulation of layer-specific neurons is disrupted in *Piwil1* KO. (A) Immunohistochemisty for markers of upper-layer neurons Cdp (red) and Satb2 (green) and lower-layer markers Tle4 (green) and Ctip2 (red) on WT and *Piwil1* KO coronal sections. Confocal images were equally subdivided into 10 bins for analysis from cortical plate (bin 1) to ventricular zone (bin 10). (A’) Analysis was performed by counting number of Cdp+ and Tle4+ neurons or Bcl11b+ and Satb2+ neurons in each bin. Quantification of the distribution of each marker is given as the percentage from total DAPI+ neurons. (B,B’) Quantification of total number of Satb2+ and Ctip2+ across bins and (C) co-localization within same cells of Satb2+ with Ctip2+.

### Computation of piRNA expression levels

The genome-mapped reads were compared to the known testicular piRNAs downloaded from the piRNAbank database (29). From the remaining genome-mapped reads that were not in the known piRNA database, a list of the top putative brain-expressed piRNAs was identified based on the presence of a first uridine (1U) nucleotide and a length ~30nt. The transcripts per million (TPM) measure for all piRNAs and putative piRNAs were then computed. This is a standard measure used in RNA-seq analysis and is more appropriate for small RNA-seq analysis than other commonly-used measures such as reads per kilobase per million reads (RPKM) because the piRNAs are all very short and essentially the same length.

We computed differential gene expression between the WT and KO samples as the log (base 2) fold change of the average TPM in the KO samples over the average in the WT samples. There are a number of published bioinformatics methods for detecting differentially expressed genes from RNA-seq data, such as the DESeq method (30). These methods generally make an assumption that the majority of genes do not change in expression between samples and therefore the median log fold change over genes can be used as a scaling factor to scale all the expression values for the genes. However, this assumption does not obviously hold in our experiment since all piRNAs could potentially change in expression in the knockout mouse strain and we do not have a high confidence set of mouse brain piRNA loci to use as a reference, comparable to protein coding genes.

We thus used a simpler approach of searching for the highest log fold changes between WT and KO without scaling, with the caveat that we may miss large global changes in piRNA expression with this approach. We note that this is a general issue with RNA-seq as a technique for measuring gene expression since RNA-seq is by nature a relative measure of transcript abundance, unlike microarrays. We also tested whether accounting for outlier samples using a geometric mean instead of an arithmetic mean had any effect (similar to the DESeq method), but did not find any major effect on our data set.

### tRNA-derived reads analysis

We downloaded a total of 433 mouse tRNA sequences from the UCSC genome browser (mm10). These tRNA genes encode 22 types of tRNAs and 52 types of tRNA isoacceptors. We computed the TPM measure for all tsRNAs for downstream analyses. The 3’ versus 5’ bias is calculated as the log 2 ratio of 3’ tsRNAs over 5’ tsRNAs (i.e., log2(3’/5’)).

## Results

### Neocortical zone specific disruption of cell cycle in *Piwil1* KO mouse

We previously reported that *Piwil1* is expressed in VZ of E17 neocortices (5). To test the effects of *Piwil1* depletion on the E17 neocortical RG cell cycle, we injected the thymidine analogue 5-chloro-2’-deoxyuridine (CldU, 50 mg/kg) at E16, 24 hours before fixation, and 5-iodo-2’-deoxyuridine (IdU, 50 mg/kg) at E17, 1 hour before fixation. The CldU labeled RG that have exited the cell cycle and IdU labeled RG in S-phase of the cell cycle (25,31). RGs that are in M-phase were immmunolabeled using phospo-Histone3 (pH3). Confocal analysis followed by double blind quantification revealed that total cell cycle re-entry across the developing neocortical wall was unaffected in *Piwil1* KOs (Fig 1A). However, when neocortex was split into 10 bins from pial to ventricular surface, we found zone specific effects on cell cycle re-entry in the developing neocortex depleted of *Piwil1* (Fig 1B). In particular, we found decreased cell cycle re-entry in the ventricular zone (VZ) enriched for RG coupled with increased cell cycle re-entry in the sub-ventricular zone (SVZ) enriched for intermediate progenitor cells (IPCs). Furthermore, we found that cycling cells in *Piwil1* KO are increased in number in their mitotic phase (M-phase) of the cell cycle, particularly in the VZ (Fig 1C,D). Finally, at postnatal day 0 (P0), we found increased amount of Pax6+ radial glia (RG) progenitors in the VZ (Fig 1E,F), and no change in Tbr2+ intermediate progenitors (not shown). These data indicate spatial-specific effects of Piwil1 on cycling with a particular Piwil1 role in progression of neural stem cells through the M-phase.

RG progenitors serve as a scaffold for neuronal migration (3). Since their cell cycle progression was disrupted in *Piwil1* KOs but we still found some of cells to exit the cell cycle, we assessed if migration of born neurons is also affected after *Piwil1* was silenced using specific shRNAs or deleted in *Piwil1* KOs. We found that *Piwil1* depletion exhibited migration defects of projection neurons in the developing neocortex similar to recent report by Zhao et al. (14; not shown). To characterize the role of *Piwil1* in neuronal migration of distinct subpopulations of neocortical projection neurons, we next examined distribution of molecular markers expressed in a layer-specific manner in *Piwil1* KO mice at postnatal day 0 (P0) and P7. We immunostained Tle4 and Ctip2 to identify typically deep-layer subcortically projecting neurons (DL), and Cdp for upper-layer (UL) intracortically projecting neurons. Additionally, we stained for Satb2 for upper and lower layer post-mitotic neurons. As reported by Zhao et al (Mol Brain 2015), we did not find changes in relative distribution of tested markers neither in primary nor secondary sensorimotor regions of *Piwil1* KOs when compared to littermate WTs (Fig 2A). However, we found an increased total number of Satb2 and Ctip2 across bins that was coupled to increased co-localization within same cells of Satb2 with Ctip2 (Fig 2B). These data suggest that Piwil1 plays a role in postmitotic identities of neocortical projection neurons. Collectively, these data indicate Piwil1 plays a role in cell cycle, neuron migration, and possibly in post-mitotic specification of distinct neuronal subpopulations.

### Prenatal *Piwil1* silencing disrupts neocortical dendritogenesis

Because we found migratory defects in *Piwil1* KO, which usually lead to disrupted neocortical circuits, we assessed whether decreased connectivity is associated with defects in dendritogenesis. We performed quantification on Golgi-stained brains, analyzing upper and lower neocortical layers from adult WT and *Piwil1* KO mice. We used Neurolucida software for 3D reconstruction and quantification in double blind fashion (Fig 3A). *Piwil1* KO animals displayed a decrease in dendritic length and spine number in upper layer neurons, while lower cortical layers were unaffected. These findings indicate that constitutive prenatal loss of *Piwil1* affects formation of cortical circuits resulting in abnormal cytoarchitecture in the postnatal and adult neocortex.

**Figure 3.**
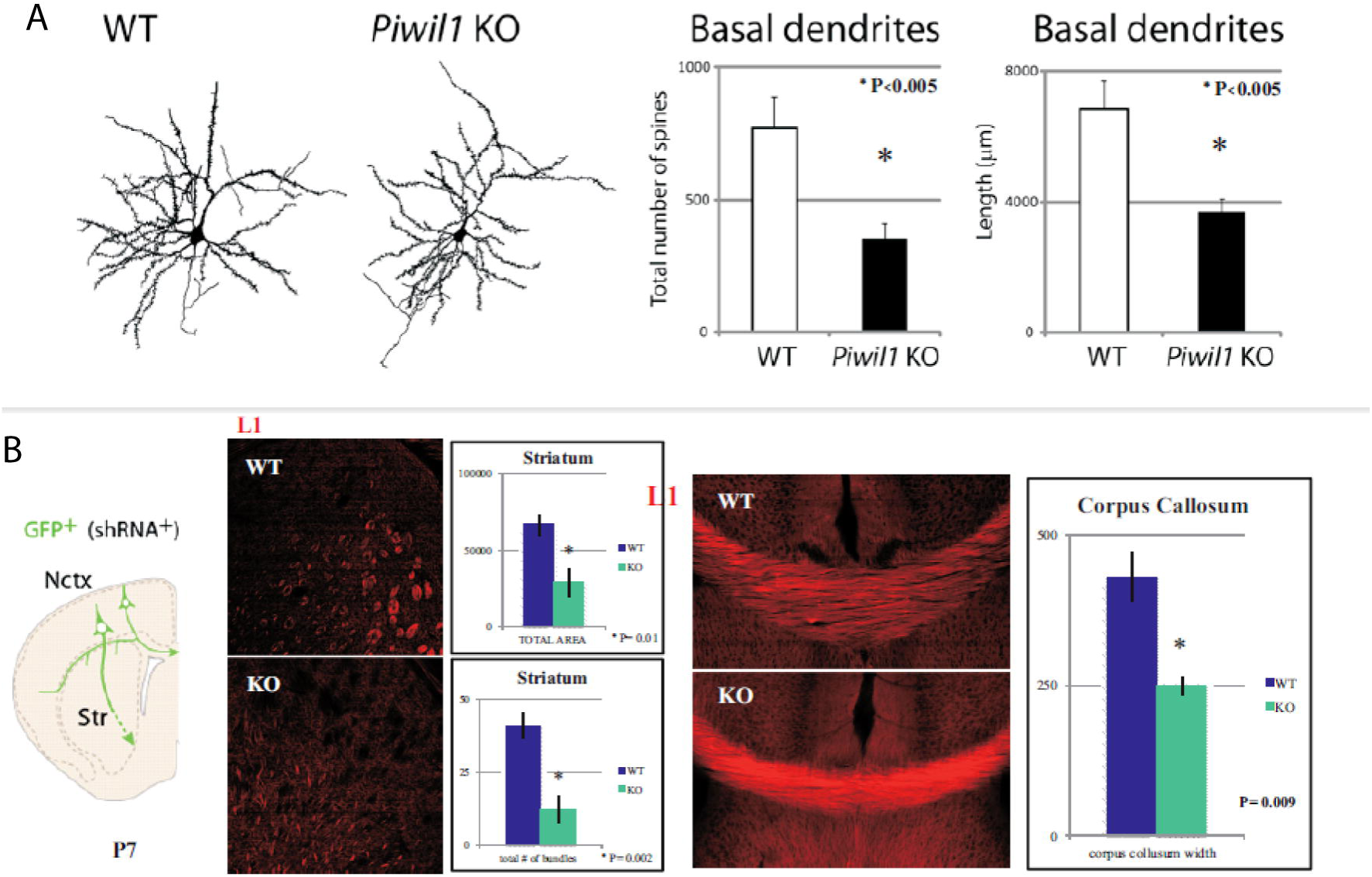
Loss of *Piwil1* results in abnormal cytoarchitecture in the adult neocortex. (A) Representative reconstructions of neocortical primary neurons of adult WT and *Piwil1* KO and quantification of length and spine number of basal dendrites. (B) Schematic of upper-layer neuron projecting intracorticaly while lower-layer is projecting subcorticaly at P7. Immunohistochemistry for axonal marker L1 of cortico-striatal bundles and corpus callosum in WT and *Piwil1* KO. Quantification of positive area and positive number of bundles and quantification of corpus callosum width showing reductions in KO.

Since we found aberrant identity of neocortical projection neurons, we next examined cortico-striatal bundles and corpus callosum features in *Piwil1* KO mice at P7 (Fig 3B). Using the axonal marker L1, we compared cortico-striatal axonal bundles based on number and size. Both these measurements showed significant reductions, suggesting disrupted cortico-striatal connections. Additionally, the width of the corpus callosum was significantly reduced, suggesting reduction in intracortical connectivity. Thus, we conclude that prenatal *Piwil1* silencing disrupts neocortical circuits, which was associated with a number of neurodevelopmental disorders.

### Sequencing piRNA in developing neocortex

Next, we investigated the expression of piRNAs in neocorticogenesis and the effect of *Piwil1* depletion on their expression. To identify piRNAs that potentially affect the phenotypes we observed, we extracted and sequenced small RNAs with known piRNA characteristics (i.e. length and sequence composition) from KO and WT neocortices. As a control we sequenced small RNAs from testes (known to express piRNAs) and spleen (which is not known to express piRNAs). When we aligned the sequencing reads to known non-coding, non-Piwi-interacting RNAs (ncRNAs) and tRNAs, ~45%-50% of reads in neocortices, ~6% of reads in WT testes, ~9% of reads in KO testes and ~40% of reads in spleen mapped to the ncRNAs or tRNAs (Table 1).

**Table 1.**
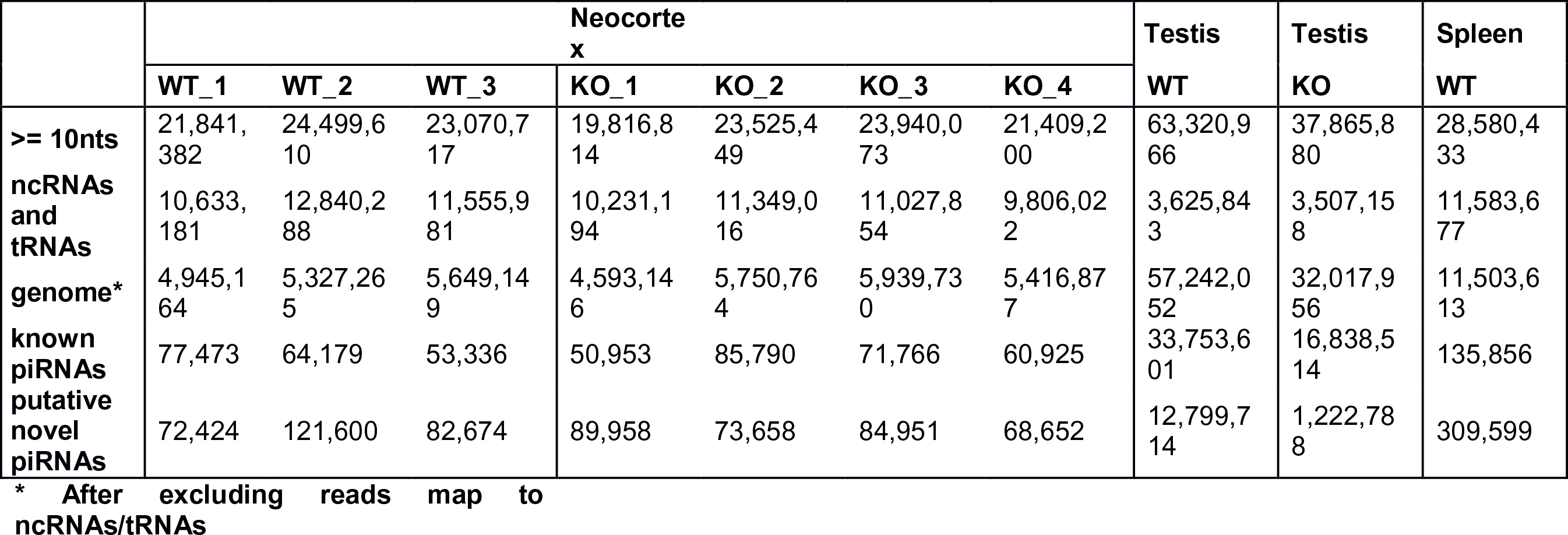
Summary of the number of reads aligned in samples.

We removed all reads aligned to ncRNAs and tRNAs and aligned the remaining reads to the mouse genome. We compared reads aligned to the mouse genome with a database of known testicular piRNAs (Table 1). In all neocortices and spleen samples, a small percentage of reads aligned to known piRNAs from mouse testes (~1%). This number was much higher in both WT and KO testes samples (59% and 53%, respectively). This result may reflect the fact that neocortical piRNAs are different than testicular piRNAs, that there are neocortical piRNAs derived from ncRNAs/tRNAs (a possibility we explore below), or that neocortical piRNAs are expressed at low levels or in only a small number of cells in our mouse neocortex samples.

Next we considered potential neocortex-specific piRNAs, which we defined as reads mapping to the mouse genome which fit two criteria: the sequence starts with one U nucleotide; and is between 29 and 31nt in length. Overall the number of potential neocortex-specific piRNAs was similar to that of known testicular piRNAs (Table 1).

We examined the highest expressed known and potential piRNAs. All of the highest expressed transcripts were all known testicular piRNAs, with expression levels roughly ten times higher than the potential neocortical piRNAs (Supplemental Table 1). This strongly suggests that our piRNA sequencing protocols and bioinformatics analysis correctly identified real piRNAs. It also suggests that neocortical piRNAs are similar to or expressed from similar loci as the testicular piRNAs, but are expressed at much lower levels or only in specific cell types in the neocortex. We cannot distinguish between these two possibilities due to the heterogeneity of our samples.

Nonetheless, we did not find either high absolute expression of piRNAs or significant differential expression of piRNAs between WT and *Piwil1* KO mice. Indeed the top-four highest expressed piRNAs were the same between WT and KO mice (Supplemental Table 1). This suggests that the phenotypes caused by *Piwil1* knockout may occur independently of piRNAs, a possibility that has previously been suggested using HITS-CLIP experiments (38).

### tRNA-derived reads in developing neocortex

Serendipitously, we found that in the neocortical samples, on average >12% of the small RNA reads mapped to tRNAs, as compared to <1% for the testis samples. Many of the tRNA-derived reads ended in CCA (~72% of brain reads), which is typically seen on the ends of mature tRNAs. This enrichment of 3’ tRNA fragments might be partly due to the protection of the 3’ end of charged tRNAs from the periodate treatment during sequencing library preparation (see methods for detail). Nevertheless, similar functional tRNA-derived small RNAs (tsRNAs) have been reported in multiple recent studies (32–34,43,44). In particular, Keam *et al.* (33) reported that the human Piwi ortholog, Hiwi, binds to tsRNAs in cancer cells, based on immunoprecipitation results. Thus our data suggests that tRNA-derived piRNAs are expressed in the developing mouse neocortex and might have a function related to mouse neocorticogenesis.

To determine the origin and properties of these tsRNAs, we looked at the tsRNA distribution among the 433 mouse tRNA genes. These tRNA genes encode 22 types of tRNAs and 52 types of tRNA isoacceptors. Over all samples, we found that the most highly expressed tsRNAs remained stable across tissues. tsRNAs derived from five tRNA isoacceptors – TyrGTA, LysTTT, LysCTT, GluCTC and GluTTC – were consistently the highest expressed across all brain samples, comprising 40%-44% of all tsRNA reads from these samples (Supplementary Table 2). Similarly, tsRNAs derived from Glu, Lys, Ala are the highest expressed at the tRNA level (Supplementary Table 3). Furthermore, there was a consistent 3’ vs. 5’ bias in the processing of the tRNAs that differed for each tsRNA but was consistent across samples (Pearson correlation 0.998 for comparisons between brain WT and KO samples, Supplementary Table 4). Such biases in tsRNA processing have been previously reported in other piRNA studies, such as Keam et al. (33) and a study in marmoset by Hirano et al. (42). We also found an overall bias toward a U in the first base in brain (69% in 3’ biased reads and 54% in 5’ biased reads), consistent with an association with piRNAs and Piwi protein.

We also found consistencies when comparing our tsRNAs to those reported from Piwi IP studies in marmoset (42) and human cancer cells (33). Of the 5 most highly expressed tsRNAs in Ref. 42 and Ref. 33, 4 and 3 respectively were among the 10 most highly expressed in our study. Furthermore, the 5’ bias of these shared tsRNAs was observed in both our study and their studies. Taken together, the non-random processing of the tRNAs into small RNAs and consistency with previous studies using different methods based on Piwi immunoprecipitation strongly suggests that the small RNAs might be functional as opposed to being random degradation products. Our results add to the literature on tRNA-derived RNAs and suggest that these may be a potentially interesting class of RNAs to study in the future in the context of neocortex development.

## Discussion

The functional roles of PIWI proteins and their associated piRNAs have been best studied in the germline of a wide range of animal species. A number of studies also studied a possible role for piRNAs in the CNS (17,18,35,36). Of these, the most detailed study is perhaps that of Rajasethupathy *et al.* (17) which elucidated the roles on piRNAs in memory formation in Aplysia. However, some lingering doubts remain among some researchers about the magnitude of the role of piRNAs in the mammalian brain because of the low expression levels of the piRNAs and concern that they may represent degradation products instead of true piRNAs. Another issue is that bioinformatic sequence matches of piRNAs to non-transposon targets such as promoter regions is difficult and not easily amenable to statistical analysis (e.g. Ref. 17). Nonetheless, we previously showed that *Piwil1* mRNA, encoding piRNA associated protein, expressed in developing neocortical wall increased from early (E13) to late neurogenesis (E15, E18) (5). In addition, an RNAi knock-down study revealed a *Piwil1* role in neocortical migration (14). We tested the role of both PIWI-like 1 protein and piRNAs in the developing mouse neocortex by ablating *Piwil1* from developing neocortical wall. We confirmed the results of Zhao *et al.* (14) that Piwil1 is critical for normal neuronal migration. In addition, we showed that during neocorticogenesis, constitutive KO of *Piwil1* affects cell cycle dynamics with compartmentalized and selective effects on M-phase of the cell cycle, migration of later born neurons, and neocortical circuitry formation, particularly disrupting the dendritogenesis of upper layer neurons. We also found that prenatal ablation of *Piwil1* resulted in an increased co-localization of Satb2 and Ctip2. Collectively, these results implicate that loss of *Piwil1* early in neocortical development results in abnormal neocorticogenesis at multiple levels, suggesting that it acts on different downstream targets at different time points during neocortical development. In summary, we conclude that prenatal *Piwil1* silencing disrupts neocortical circuits, which was associated with a number of neurodevelopmental disorders. Interestingly, PIWI mutations were recently associated with neurodevelopmental autism spectrum disorders (15).

*Piwil1* KO-associated shortening of dendrites and spine reduction is restricted to basal and not apical dendrites, which suggests that the Piwil1 silencing complex has selective roles within dendritic compartments of single neurons. However, this can also be secondary consequence of disrupted axonal projections from the contralateral side. Additionally, the activity of Piwil1 at one developmental stage in upper layers may be substantially different than its activity in another due to differential temporal expression of RNAs available to act upon. There are numerous potential stimuli for dendrite lengthening and the subset of stimuli for which Piwil1 plays a part in, and may contribute to neurodevelopmental disorders associated with disrupted neocortical circuitry development, such as autism.

In the course of sequencing piRNAs in the developing neocortex, we serendipitously discovered a large class of piRNAs derived from tRNAs. This class of piRNAs has commonly been discarded in bioinformatic analysis of piRNAs because it is assumed to represent degradation products. In contrast we showed that these piRNAs are non-randomly processed from tRNAs and therefore potentially biologically functional. A functional role for tRNA-derived piRNAs was previously demonstrated by several studies, including a PIWI immunoprecipitation study in human cancer cells (33), another study that demonstrated a role for a tRNA-derived RNA in cell proliferation (37) and two recent studies demonstrated a role for tRNA-derived RNAs in mammalian sperm. Thus our results provide further evidence that this class of RNAs may be interesting objects of future study.

Overall, our data helps clarify the somewhat conflicting reports on Piwi and piRNA activity in the brain. Although our experiments show that Piwil1 has an important role in mouse neocorticogenesis, our sequencing experiments revealed relatively low expression for piRNAs in the neocortex. Previous research also showed a wide range but generally low expression for piRNAs in mouse hippocampus (18). While we cannot exclude the possibility that piRNAs with low-level expression may nonetheless play a significant biological role, we do not find statistically significant evidence of differential expression of piRNAs between WT and KO mice. Such a piRNA-independent function is supported by a previous study (38) that used Piwil1 HITS-CLIP to show that Piwil1 binds spermiogenic mRNAs directly without a piRNA intermediate. Similar results have also been shown by the RNA immunoprecipitation experiments in Zhao *et al.* (14). Similarly, a recent study reports that Argonaute proteins have sequence specificity in the absence of microRNAs (39). In addition, Piwil1 regulates mRNA translation (40), one of key regulatory mechanisms in neocortical development (5,12). Finally, in our own data, all the top differentially expressed piRNAs were previously known piRNAs from testes, as opposed to putative neocortex-specific piRNAs. This strongly suggests that our piRNA sequencing results are accurate and not simply noise. Taken together, these results suggest that Piwil1 may have a piRNA-independent function in mouse neocortices that contribute to multiple steps of neocortical development.

## Acknowledgements

This work was supported by National Institutes of Health (NIH) grants (NS064303 and NS075367) and Robert Wood Johnson Medical School start-up funds to M.R.R. M.K. was supported by a grant from the NIH (GM067005). We thank Drs. Althea Stillman and Jack Pike for technical support.

---

Supplementary figure 1: The size distribution of sequencing reads for a representative sample (3WT).

Supplementary figure2: ExampleofatRNAalignmentwitha3-nt“buffer”:

